# Lightweight Fine-tuning a Pretrained Protein Language Model for Protein Secondary Structure Prediction

**DOI:** 10.1101/2023.03.22.530066

**Authors:** Wei Yang, Chun Liu, Zheng Li

**Affiliations:** Henan Key Laboratory of Big Data Analysis and Processing,Henan Engineering Laboratory of Spatial Information Processing, School of Computer and Information Engineering,Henan University, Kaifeng, 475004, China

**Keywords:** Protein language model, Parameter efficient fine-tuning, Protein secondary structure prediction, Nine-class prediction

## Abstract

Pretrained large-scale protein language models, such as ESM-1b and ProtTrans, are becoming the fundamental infrastructure for various protein-related biological modeling tasks. Existing works use mainly pretrained protein language models in feature extraction. However, the knowledge contained in the embedding features directly extracted from a pretrained model is task-agnostic. To obtain task-specific feature representations, a reasonable approach is to fine-tune a pretrained model based on labeled datasets from downstream tasks. To this end, we investigate the fine-tuning of a given pretrained protein language model for protein secondary structure prediction tasks. Specifically, we propose a novel end-to-end protein secondary structure prediction framework involving the lightweight fine-tuning of a pretrained model. The framework first introduces a few new parameters for each transformer block in the pretrained model, then updates only the newly introduced parameters, and then keeps the original pretrained parameters fixed during training. Extensive experiments on seven test sets, namely, CASP12, CASP13, CASP14, CB433, CB634, TEST2016, and TEST2018, show that the proposed framework outperforms existing predictors and achieves new state-of-the-art prediction performance. Furthermore, we also experimentally demonstrate that lightweight fine-tuning significantly outperforms full model fine-tuning and feature extraction in enabling models to predict secondary structures. Further analysis indicates that only a few top transformer blocks need to introduce new parameters, while skipping many lower transformer blocks has little impact on the prediction accuracy of secondary structures.

## 1. Introduction

Pretrained large-scale protein language models, such as ESM-1b (Rives et al., 2021) and ProtTrans (Elnaggar et al., 2021), are increasingly popular for biological modeling based on amino acid sequence information. The embedding features extracted from these models have been shown to work well for various downstream tasks, including protein secondary structure prediction (Høie et al., 2022; Singh et al., 2022b; Yang et al., 2022b), contact prediction (Singh et al., 2022a), subcellular localization (Stärk et al., 2021), protein succinylation site prediction (Pokharel et al., 2022), fold recognition (Villegas-Morcillo et al., 2022), and variant effect prediction(Marquet et al., 2022). In particular, compared with traditional profile features, such as PSSM and HHM, which are obtained by using PSI-BLAST (Altschul et al., 1997) or Hhblits (Remmert et al., 2012) to perform time-consuming multiple sequence alignment operations against protein sequence databases, embedding features not only allow fast inference but also perform better on multiple tasks. Recently, ESM-2 (Lin et al., 2022) further increased the number of parameters of the transformer model to 15 billion and experimentally demonstrated that larger models can provide better embedding representations for protein structure prediction. In addition, the number of protein sequences that can be used to train a transformer model grows exponentially. As a result, the size of protein language models will become increasingly large to better capture the biological information implied in protein sequence data.

Protein language models are obtained by training transformer models on massive unlabeled protein sequence data in a semisupervised manner. The goal of the training is to predict masked residues according to the input amino acid sequences with partially masked residues. Therefore, the knowledge learned by pretrained protein language models is general and task-agnostic. For downstream tasks, both feature extraction and fine-tuning can be used. In feature extraction, the weights of the pretrained model are fixed and the extracted embedding representations are used as input features for a downstream model. For fine-tuning, the parameters of the pretrained model and the downstream model need to be tuned simultaneously according to a given target dataset. In fact, fine-tuning pretrained models has become a standard paradigm in natural language processing (Devlin et al., 2018; Liu et al., 2019; Raffel et al., 2020) and computer vision (He et al.; Li et al.). Many studies have shown that fine-tuning can enable models to learn task-specific representations and thus, compared to feature extraction, can lead to substantial performance gains (Dodge et al., 2020; Howard and Ruder). Moreover, the literature (Kumar et al.) also found that fine-tuning performs worse in the presence of a large distribution shift. However, for protein-related modeling tasks, recent works (Høie et al., 2022; Yang et al., 2022b) are all based on feature extraction. Whether fine-tuning is superior to feature extraction on these tasks has not been specifically investigated. The primary sequences of all proteins consist of 21 amino acid abbreviation letters (which correspond to 20 standard amino acids and 1 nonstandard amino acid). Therefore, it is unlikely that the sequence data used in the pretraining of protein transformers and the sequence data used in the transformers’ downstream tasks will have large distribution shifts. Therefore, we believe that the judicious use of fine-tuning can improve the performance of the protein transformers’ downstream tasks. In particular, considering that protein secondary structure prediction is a widely studied fundamental problem in computational molecular biology, we focus only on such prediction in this paper. We expect that the findings in this study can inspire research on other downstream tasks.

The most common way to fine-tune a pretrained model is to update all the model parameters. However, for large models, this full fine-tuning is prohibitively expensive in terms of computation and storage. This issue becomes more salient as the size of the pretrained model continues to increase. To alleviate this issue, several lightweight fine-tuning methods, such as Adapter (Houlsby et al., 2019), Compacter (Mahabadi et al.), Prefix-tuning (Li and Liang), LoRA (Hu et al.), (IA)^3^ (Liu et al., 2022), MAM adapter (He et al., 2022), and UNIPELT (Mao et al.), have been proposed; these methods freeze the original pretrained parameters and tune only a few newly introduced parameters in each transformer block of the pretrained model. In particular, the newly introduced parameters allow frozen pretrained models to learn task-specific representations and avoid storing large checkpoints during training. This parameter-efficient tuning also significantly reduces the computational and memory overhead per iteration since the gradients of the pretrained parameters do not need to be computed and stored during backpropagation. More importantly, lightweight fine-tuning has performed competitively against full fine-tuning on a variety of tasks. Therefore, in this paper, we employ lightweight fine-tuning techniques to develop a novel framework for predicting protein secondary structures.

Secondary structures (SS) are locally folded structures formed within a polypeptide and are important for analyzing the relationship between protein structures and amino acid sequences. To obtain an unambiguous and physically meaningful definition of SS, Kabsch and Sander (Kabsch and Sander, 1983) designed a program, DSSP, which identifies hydrogen bonding patterns and geometric features from the atomic coordinates of a protein with a known structure and then assigns a secondary structure type to each amino acid residue in the protein according to some criteria. In (Kabsch and Sander, 1983), secondary structures were classified into eight categories: *α*-helix (H), *β*-bridge (B), strand (E), 3_10_-helix (G), *π*-helix (I), turn (T), bend (S), and loop (L). These eight categories are often further reduced into three categories: helix (H, G, and I), strand (B and E), and coil (T, S and L). The DSSP program was rewritten in 2015 to better identify *π*-helices (Touw et al., 2015). The latest version, DSSP 4.0, was released in August 2021. Currently, the .dssp files in the DSSP database are all generated based on DSSP 4.0. In particular, the biggest change in DSSP 4.0 compared to the previous version is the expansion of the original eight secondary structure types to nine, i.e., the addition of a new helix type, the poly-proline helix (P). DSSP is the de facto standard used by the PDB database to assign secondary structures. Therefore, for a protein whose structure is known, the secondary structure sequence output by the latest DSSP according to its structure file is considered the ground truth label information.

For a given protein chain, the goal of secondary structure prediction is to assign a secondary structure type to each residue in the chain according to only its amino acid sequence information. Protein secondary structure prediction is clearly a classification problem. The earliest work on protein secondary structure prediction can be traced to 1976 (Levitt and Chothia, 1976). Since then, a variety of neural network-based secondary structure predictors, such as Porter (Baldi et al., 1999), PSIPRED (Jones, 1999; McGuffin et al., 2000), SSpro (Pollastri et al., 2002), Jpred3 (Cole et al., 2008), and MBRNN (Babaei et al., 2012), have been proposed. These predictors are aimed mainly at simple three-class secondary structure prediction because the known-structure protein data available for training were extremely limited at that time. In particular, compared with coarse-grained three-class prediction, fine-grained eight-class prediction can provide more detailed local structure information. Therefore, subsequent work focuses on improving the prediction accuracy of the eight-class secondary structure. For example, RaptorX-SS8 (Wang et al., 2011) uses conditional neural fields for eight-class secondary structure prediction. To learn hierarchical representation for prediction, Zhou and Troyanskaya (Zhou and Troyanskaya) developed a supervised generative random network. DCRNN (Li and Yu) exploits a cascaded multiscale convolutional network and bidirectional gate recurrent units to extract local and global contextual features. DeepCNN (Busia and Jaitly, 2017) has a novel chained convolutional architecture and uses next-step conditioning to improve performance on eight-class secondary structure prediction. Drori et al. (Drori et al., 2018) investigated the performance of eight-class prediction under six different deep network architectures. MUFOLD-SS (Fang et al., 2018) uses a deep inception-inside-inception network and hybrid profile features for prediction and achieves state-of-the-art performance. To effectively capture both local and nonlocal interactions between amino acid residues, DeepACLSTM (Guo et al., 2019) proposes a deep asymmetric convolutional long short-term memory neural model. Following this idea, SAINT (Uddin et al., 2020) further introduces a self-attention mechanism for the deep inception-inside-inception network. Moreover, multitask learning-based deep predictors, such as NetSurfP-2.0 (Klausen et al., 2019), SPOT-1D (Hanson et al., 2019), and OPUS-TASS (Xu et al., 2020), have also been proposed. Recently, ShuffleNet_SS (Yang et al., 2022a) demonstrated that state-of-the-art prediction performance can also be achieved by using a lightweight convolutional network with few parameters. To predict protein seconary structure, Protein encoder (Uzma et al., 2023) first utilizes an unsupervised autoencoder to perform feature extraction, and then uses ensemble of three feature selection methods to select the optimum feature subset. The input features used by the predictors mentioned thus far are mainly profile features derived from multiple sequence alignments. However, performing multiple sequence alignment operations against ever-growing protein sequence databases is extremely time-consuming. Consequently, recent work has employed mainly embedding features, which are extracted from pretrained protein language models, to develop secondary structure prediction methods. Representative predictors include DML_SS (Yang et al., 2022b), SPOT-1D-LM (Singh et al., 2022b), and NetSurfP-3.0 (Høie et al., 2022). A more detailed review on protein secondary structure prediction can be found in (Ismi et al., 2022).

For all existing secondary structure predictors, the secondary structure information used in their training and evaluation is derived from the eight-class secondary structure assignment of the DSSP program. Although recent works, such as DML_SS (Yang et al., 2022b), NetSurfP-3.0 (Høie et al., 2022), and SPOT-1D-LM (Singh et al., 2022b), were published after the release of DSSP 4.0, they still do not use the secondary structure information provided by the nine-class secondary structure assignment, thereby implying that they are trained and evaluated on outdated label information. To this end, in this study, we will use the latest secondary structure sequence information for training and evaluation. To the best of our knowledge, this is the first work developed for nine-class secondary structure prediction.

In addition, for protein secondary structure prediction, to ensure nonhomology between the training and test sets, it is usually required that all protein chains in the training set have no >25% sequence identity with any protein chain in the test set. Influenced by this convention, existing works, such as DML_SS (Yang et al., 2022b), NetSurfP-3.0 (Høie et al., 2022), RaptorX-SS8 (Wang et al., 2011), SPOT-1D (Hanson et al., 2019), and OPUS-TASS (Xu et al., 2020), also typically require no more than 25% or 30% sequence identity between any two protein chains in the training set. A lower sequence identity cutoff means that more protein chains with known structures cannot be used as training data. However, for deep models, more training data usually mean better predictive performance. In particular, increasing this cutoff value does not violate the nonhomologous convention between the training and test sets. Therefore, to take full advantage of protein data with known structures, the training set should be constructed based on a higher sequence identity cutoff. To determine the appropriate cutoff value, we investigate the impact of training sets constructed based on different cutoff values on the performance of secondary structure prediction. Protein secondary structure prediction has been conducted for more than 40 years; however, we found that no work has yet been done to carry out such investigations.

In this paper, we propose a novel end-to-end protein secondary structure prediction framework that involves the lightweight fine-tuning of a pretrained protein language model. The proposed framework consists mainly of a pretrained transformer model and a prediction head. For the pretrained transformer model, we retain its original parameters as fixed and experiment with seven state-of-the-art parameter-efficient finetuning methods to inject new parameters into it. The prediction head is a convolutional network with a multibranch topology that uses layer normalization instead of batch normalization, thus eliminating the effect of PAD tokens on non-PAD tokens. In particular, we need to update only the newly introduced parameters and parameters of the prediction head during training, thereby significantly reducing memory overhead. Moreover, we investigate the impact of softmax-based metric learning loss and cross-entropy classification loss on the performance of protein secondary structure prediction.

In summary, the contributions of this work are as follows. (1) We propose a novel lightweight fine-tuning framework for protein secondary structure prediction. To our knowledge, this is the first empirical study of the lightweight fine-tuning of a pretrained protein language model for a protein-related biological modeling task. (2) Extensive experiments on seven test sets show that the proposed methods outperform existing state-of-the-art predictors. (3) We experimentally demonstrate that lightweight fine-tuning outperforms feature extraction and full model fine-tuning on both three-class and nine-class predictions. In particular, this study is the first attempt to perform nine-class secondary structure prediction. (4) We comprehensively evaluate the performance of seven state-of-the-art lightweight fine-tuning methods (Adapter, Compacter, Prefix-tuning, LoRA, (IA)^3^, MAM adapter, and UNIPELT) on protein secondary structure prediction. The results demonstrate that only a few top transformer blocks need to introduce new parameters, while skipping many lower layers has little impact on the prediction accuracy of secondary structures. (5) We experimentally show that simply increasing the sequence identity cutoff between protein chains can increase the size of the training data and thus significantly improve the prediction accuracy of secondary structures.

The remainder of the paper is organized as follows. Section 2 introduces the proposed protein secondary structure prediction framework in detail. In Section 3, the datasets used, implementation details and experimental results are given. We conclude this paper in Section 4.

## 2. The proposed method

In this section, we propose an end-to-end framework for protein secondary structure prediction, as shown in Fig.1. The framework consists of four modules: the input, output, pretrained transformer model, and prediction head. The input is an amino acid sequence that consists of 21 amino acid abbreviation letters {A, C, D, E, F, G, H, I, K, L, M, N, P, Q, R, S, T, V, W, Y, and X}, and the output is a protein secondary structure sequence, which is obtained by assigning a secondary structure type to each amino acid residue in the input. For batch processing, it is usually necessary to add two special tokens, namely, EOS and PAD, to the right side of the amino acid sequence, where the former represents the end of a protein chain and the latter is used to make all amino acid sequences in a minibatch the same length. In particular, the PAD tokens will be ignored by the attention mechanism in the pretrained transformer model, which is obtained by self-supervised learning on a large-scale protein sequence dataset. Existing pretrained transformer models typically contain hundreds of millions of parameters, and the size of these models will continue to increase, so updating all their parameters directly requires more storage space. To this end, this study adopts lightweight fine-tuning strategies to train the proposed secondary structure prediction model. Let Θ denote the weights of a pretrained transformer model. To perform lightweight fine-tuning, some new lightweight modules need to be injected into the pretrained transformer model. The purpose of injecting new modules is to enable the pretrained transformer model to learn to encode task-specific representations while keeping Θ fixed. Therefore, only the newly introduced parameters and the parameters of the prediction head must be updated during training. In the following, we will first introduce the adopted pretrained transformer model and several lightweight fine-tuning methods, then describe the prediction head, and finally give the loss function used for model training.

**Fig. 1.**
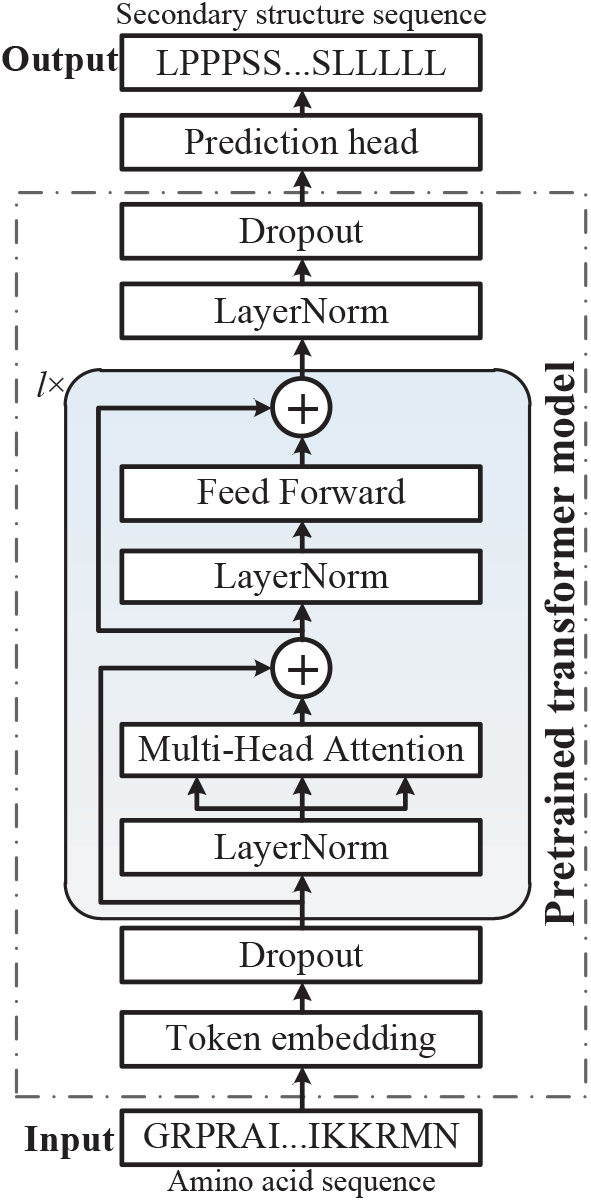
The framework for protein secondary structure prediction.

### 2.1. The pretrained transformer model

For the proposed secondary structure prediction framework, many existing protein language models, such as ESM-1b (Rives et al., 2021), ProtBert (Elnaggar et al., 2021), ProtAlbert (El-naggar et al., 2021), ProtXLNet (Elnaggar et al., 2021), ProtT5-XL-U50 (Elnaggar et al., 2021), and ProtT5-XXL-U5(Elnaggar et al., 2021), can be used as backbone networks to generate embedding features with biophysical properties for a given protein chain. Considering that the features generated by the encoder of ProtT5-XL-U50 can attain the best secondary structure prediction performance (Elnaggar et al., 2021), we adopt this model as our pretrained backbone model. In particular, ProtT5-XL-U50 is an encoder-decoder transformer model based on T5-3B (Raffel et al., 2020), and the network architecture of its encoder is also given in Fig.1. The encoder consists mainly of a token embedding and a stack of *l* = 24 identical transformer blocks. Token embedding is used to convert each token in the input sequence into a learnable numeric vector with the specified dimension *d_m_* = 1024. In particular, dropout is applied on the output of the token embedding to alleviate overfitting. Moreover, each transformer block consists of two primary modules: a multihead self-attention module followed by a feedforward module. Layer normalization and residual connection are used around each module. Specifically, let 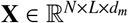 be the input tensor of a transformer block and 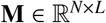 be its corresponding binary mask matrix, where *N* is the batch size, *L* is the sequence length, and *M_il_* = 0 indicates that the *l*th position in the *i*th sequence is a PAD token. The operation of two modules can be formalized as follows:

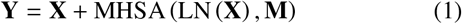

and

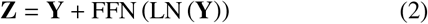

where LN denotes a simplified version of layer normalization with only rescaling and no bias term, and the functions MHSA and FFN are the multihead self-attention and the positionwise fully connected feedforward network, respectively.

In the MHSA, the input feature tensor is first transformed into three different tensors, i.e., the query tensor 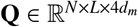, the key tensor 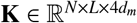 and the value tensor 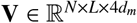, by three projection matrices, i.e., 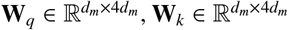 and 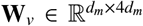, respectively. To introduce multiple heads, the shapes of the three tensors **Q**, **K** and **V** are further converted into (*N*, *h*, *L*, *d_h_*) by performing reshape and transpose operations, where *h* is the number of heads and *d_h_* = 4*d_m_/h* is the feature dimension of each head. Based on the batched matrix multiply, the attention score tensor 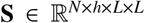 can be computed by:

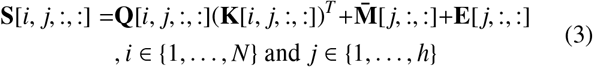

where the tensor 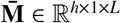 satisfies the constraint 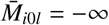 if *M_il_* = 0; otherwise, the tensor 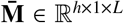 satisfies the constraint 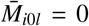; additionally, the tensor 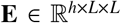 is a relative position embedding based on the key-query offsets. Different attention heads have different position embeddings, each of which is just a learnable scalar. In particular, the position embedding parameters are shared among all transformer blocks. Then, the attention weight tensor 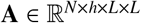 is given by:

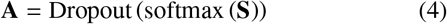

where the function softmax is applied to the last dimension of **S**, and Dropout represents a dropout layer. The attention output tensor 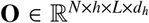 is obtained by performing the following batched matrix multiplication:

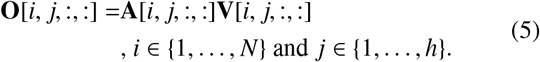

To concatenate the output of all attention heads, the attention output tensor is further transposed and reshaped, thus resulting in its shape being converted into (*N, L*, 4*d_m_).* Subsequently, the converted attention output is projected to the *d_m_*-dimensional representation space through a parameter matrix 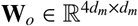. Finally, the output of the MHSA is obtained by applying another dropout.

The FFN consists of two linear transformation layers, a ReLU activation function and two dropout layers, which can be expressed as:

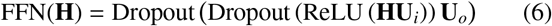

where 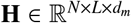 is the output of the previous layer normalization and 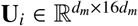 and 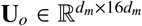 are the parameter matrices of two linear transformations. The FFN performs the same operation on the vectors at each position in the sequence.

In addition, for the final output of the stacked transformer blocks, another layer normalization is applied, followed by a dropout layer.

### 2.2. Lightweight fine-tuning methods

To perform lightweight fine-tuning on the aforementioned pretrained model, we adopt the following seven methods to inject new parameters.

#### Adapter

The adapter method (Houlsby et al., 2019) inserts a small network module called an adapter into each transformer block. In fact, the adapter can be combined with a multihead self-attention module or a feedforward module in a transformer block. In this study, we consider only the latter combination, as shown in Fig.2. There are two configuration modes for the adapter (He et al., 2022): the sequential and parallel modes. In the sequential mode, the output of the feedforward network is the input of the adapter. In the parallel mode, the input of the adapter is identical to the residual connection, which is the output of the previous multihead self-attention module. In particular, our adapter unit consists of a layer normalization, followed by a fully connected downprojection layer (FC_down_), a ReLU nonlinear activation layer, a fully connected upprojection layer (FC_up_) and a dropout layer. The FC_down_ projects the original *d_m_*-dimensional features into a lower bottleneck dimension *d*_bottieneck_, while the FC_up_ projects the features back into the original dimension. The parameter matrices of FC_down_ and FC_up_ are 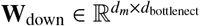 and 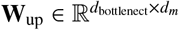, respectively. In addition, the dropout ratio in the adapter unit is set to 0.1, which is the same setting as that of the dropout layers in the pretrained model.

**Fig. 2.**
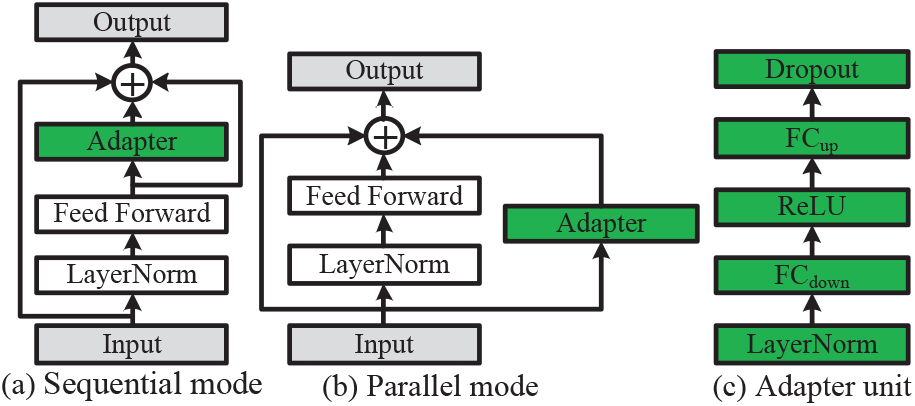
Illustration of the adapter method, which has two configuration modes: the sequential and parallel modes. (a) Sequential mode. (b) Parallel mode. (c) The architecture of our adapter unit.

#### Compacter

The compacter (Mahabadi et al.) is a parameterefficient version of the adapter; the parameter matrices in the compacter are the sum of Kronecker products between shared “slow” matrices and “fast” rank-one matrices. Specifically, let 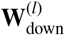 and 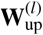 be the projection matrices of the compacter in the *l*th transformer block. Based on the Kronecker product, they are computed as follows:

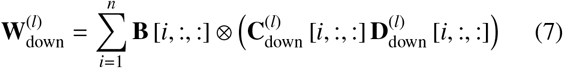

and

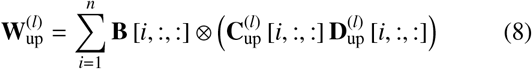

where the symbol ⊗ denotes the Kronecker product; the tensor 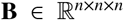 is shared across all transformer blocks in a pretrained model; and the four tensors 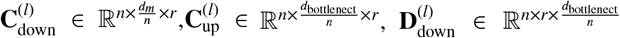 and 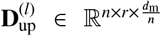 are adapter-specific parameters. Additionally, *d_m_* and *d*_bottleneck_ must be divisible by the hyperparameter *n*. In general, the compacter sets *r* to 1 and *n* to 4. We use this default setting in this study.

#### Prefix-tuning

The prefix-tuning method (Li and Liang) is used mainly in multihead self-attention, which prepends *L_p_* trainable prefix vectors to the keys and values of each head. Suppose the query tensor **Q**, the key tensor **K** and the value tensor **V** all have the shape (*N*, *h*, *L*, *d_h_*). Let 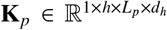 and 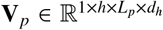 be the prefix tensors corresponding to the key and value, respectively. Then, the attention score tensor 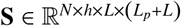 is computed as follows:

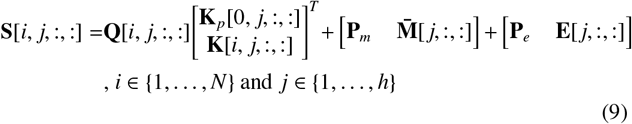

where 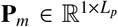 and 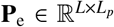 are the padding matrices with all entries being zero because each prefix vector can be regarded as corresponding to a virtual token, which does not need to be masked or to compute relative position embeddings. The attention weight tensor **A** is still computed by Eq.4, but **A**’s shape becomes (*N, h, L, Lp* + *L*). Moreover, the computation of the attention output tensor **O** in Eq.5 becomes:

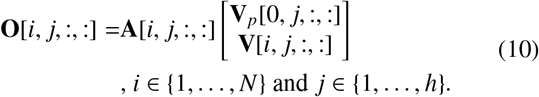

The introduction of the prefix tensors does not change the shape of the attention output tensor. In this study, we use random Gaussian initialization for the prefix tensor **K**_*p*_ and zero initialization for the prefix tensor **V***p*.

#### LoRA

Low-rank adaptation (LoRA) (Hu et al.) assumes that the changes in the weight matrices during fine-tuning of a pretrained model have a low “intrinsic rank”. For a given linear transformation, LoRA represents the update of its weight matrix as **W** + *α**W**_L_**W**^R^*, where 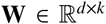 is a pretrained weight that is frozen during fine-tuning, 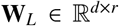 and 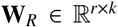 are trainable low-rank matrices, and *α* is an adjustable scalar hyperparameter with rank *r* ≪ min(*d*, *k*). LoRA applies this reparameterization to the query and value projection matrices (**W**_*q*_, **W**_*v*_) in the multihead self-attention. In particular, since **W** = **W** + *α***W**_*L*_**W**_*R*_ can be explicitly computed and stored after fine-tuning, LoRA does not incur inference latency in the testing phase. In our experiment, we set *α* to 1 and use Kaiming initialization for **W**_*R*_ and zero for **W**_*L*_.

**(IA)**^3^. The recently proposed fine-tuning method (IA)^3^ (Liu et al., 2022), which stands for “Infused Adapter by Inhibiting and Amplifying Inner Activations”, introduces three learned tensors, namely, 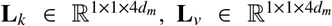 and 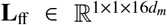, to directly modify the inner activations of each transformer block. Specifically, the first two tensors **L**_*k*_ and **L**_*v*_ are introduced to the multihead self-attention module. Let **K** and **V** be the key and value tensors, respectively, obtained by the MHSA after performing linear projections on its input features. Both **K** and **V** are of size (*N*, *L*, 4*d_m_*) at this point. Then, (IA)^3^ updates them by performing elementwise rescaling as follows: **K** = **L**_*k*_ ʘ **K** and **V** = **L**_*v*_ ʘ **V**, where the symbol ʘ denotes elementwise multiplication. Here, the broadcast mechanism of the tensor is used. Moreover, (IA)^3^ uses **L**_ff_ to modify the FFN as:

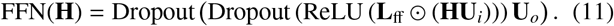

Therefore, (IA)^3^ adds only 24*d_m_* new parameters for each transformer block. In particular, to avoid changing the function represented by the pretrained model at the beginning of finetuning, (IA)^3^ initializes all of these new parameters to 1.

#### MAM adapter

The MAM adapter (He et al., 2022) is a hybrid method that simply combines parallel adapter and prefix-tuning for joint training. In (He et al., 2022), He et al. investigated various combinations of adapters, prefix-tuning and LoRA, and the results showed that the MAM adapter performed the best on natural language processing tasks. Compacter is a parameter-efficient version of the adapter. Therefore, in this study, we replace Adapter with Compacter to construct the MAM adapter.

#### UNIPELT

UNIPELT (Mao et al.) is also a hybrid method that incorporates different fine-tuning methods as submodules and automatically learns to activate suitable submodules through a gating mechanism for a specific task. Concretely, UNIPELT integrates three methods, namely, Adapter, Prefix-tuning and LoRA, into a transformer block. For each submodule, UNIPELT introduces a trainable gate 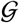 that consists of a fully connected layer (FC) with a parameter matrix 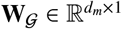 and a sigmoid activation 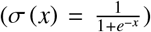. The input of the FC is the input of the module where the submodule is located (the feedforward module or multihead self-attention module). For example, in the adapter submodule, the gate output is given by 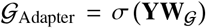, where the tensor 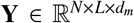 is the input of the feedforward module. Let 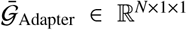 be the arithmetic mean of 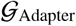 along the second axis. Then, UNIPELT uses 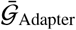 to scale the output of the adapter unit: 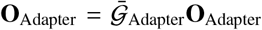. Similarly, in the prefix-tuning submodule, UNIPELT uses 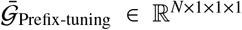 to scale the prefix tensors **K**_*p*_ and **V**_*p*_: 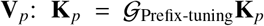, and 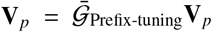. For the LoRA submodule, UNIPELT modifies the output of the linear transformation with **X**_in_ as input to: 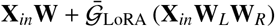. Although both 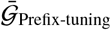 and 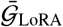 are computed based on the input of the multihead selfattention module, they employ different fully connected layers. Moreover, similar to the MAM adapter, we also use Compacter to replace Adapter in this method.

### 2.3. Prediction head

In this study, we use the embedding network proposed in (Yang et al., 2022b) as the prediction head for protein secondary structure prediction. For the sake of completeness, the network architecture of the prediction head is shown in Fig.3. As in DML_SS^embed^ (Yang et al., 2022b), the prediction head contains two inception units. Specifically, the inception unit is a two-branch network module. One branch contains two stacked basic building blocks and the other branch contains one basic building block. The output is a depth concatenation of two branches. The basic building block of DML_SS^embed^ contains batch normalization (BN). However, as pointed out in (Yang et al., 2022a), the introduction of BN can make the PAD token affect the feature generation of other non-PAD tokens during forward propagation and consequently cause inconsistent prediction results for the same amino acid sequence under different batch sizes. To this end, in this study, we replace BN with layer normalization. Fig.4 shows the new basic building block. As shown in Fig.4, we add two residual connections in the basic building block, where the dashed line indicates that the residual connection is added only when the input feature tensor and the output tensor of the layer normalization have the same shape. In particular, for the input feature tensor of the Conv1D with a kernel size of 3, we set all the features corresponding to PAD tokens as zero vectors based on the mask matrix **M**. This completely eliminates the effect of PAD tokens on non-PAD tokens. Moreover, the shape of the input feature of the prediction head is (*N*, *dm*, *L*), thus indicating that the output feature of the pretrained model needs to be transposed before being fed to the prediction head. In short, through the prediction head, each amino acid residue in an input sequence will be projected onto a 32-dimensional unit sphere 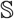^31^.

**Fig. 3.**
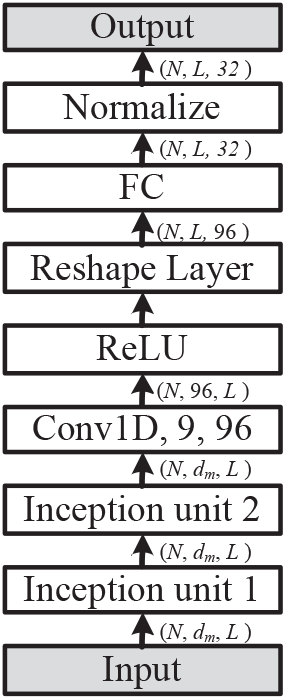
The network architecture of the prediction head.

**Fig. 4.**
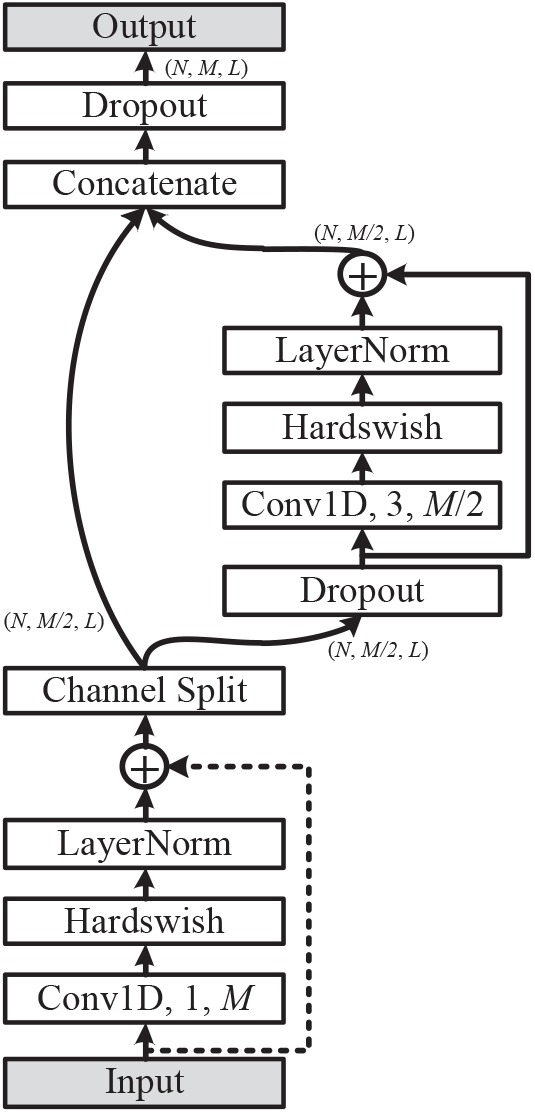
The new basic building block. Conv1D:1D convolutional layer

### 2.4. The loss function

As in (Yang et al., 2022b), we use a softmax loss based on metric learning to train the proposed secondary structure prediction model. In the embedding space, we assume that each secondary structure category has a learnable centroid. The goal of metric learning is to maximize the similarity between each amino acid residue and its target centroid and minimize the similarity between it and all nontarget centroids. Concretely, let 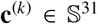 be the centroid of the *k*th secondary structure category, where *k* ∈ [0,1,…, *c* - 1] and *c* is the number of categories of secondary structures. Let 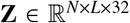 be the output tensor of the prediction head and its corresponding label matrix be 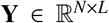, where *Y_ij_* ∈ [−100,0,1,…,*c* - 1]. The label value −100 is usually assigned to two special tokens: EOS and PAD. Then, the softmax loss can be formulated as follows:

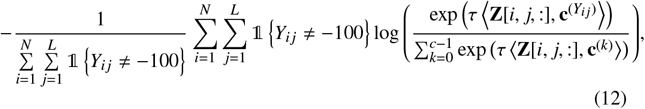

where the scaling parameter *τ* is set to 18 in this study and the indicator function 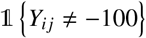 outputs 1 if *Y_ij_* ≠ −100; otherwise, it outputs 0.

## 3. Experiments

In this section, we first introduce the datasets used, then describe our implementation details, and finally report the experimental results under the different settings.

### 3.1. Datasets

In this study, we constructed protein secondary structure datasets based on the nine-class secondary structure assigned by DSSP 4.0 ^1^, released in August 2021. The original DSSP program was designed by Kabsch and Sander in 1983 (Kabsch and Sander, 1983). For each protein with a known structure, the DSSP program will generate a .dssp file containing secondary structure information based on its 3D structure data in the PDB database (Burley et al., 2020), and the resulting file is stored in the DSSP database. Currently, the .dssp files in the FTP site^2^ of the DSSP database are generated using DSSP 4.0. Compared with the previous versions of DSSP, a major feature of DSSP 4.0 is the introduction of a new secondary structure type, namely, poly-proline helix (P), on the basis of the eight secondary structure types, namely, *α*-helix (H), *β*-bridge (B), strand (E), 3_10_-helix (G), *π*-helix (I), turn (T), bend (S) and loop (L). In coarse-grained 3-state prediction, we classify the nine secondary structure types into three categories: H, G, I and P are classified as helix (H); B and E are classified as strand (E); and the other three types are classified as coil (C).

In our previous works (Yang et al., 2022a,b), we extracted primary sequence and secondary structure sequence information from the secondary structure assignment file ss.txt according to the given PDB ID and chain ID. However, we found that the file ss.txt^3^ has not been updated since August 2020. In addition, for residues whose secondary structure type is not assigned in the corresponding dssp file due to the lack of relevant atomic coordinate information in the 3D structure data, the file ss.txt uniformly sets their secondary structure type to loop (L). This actually provides incorrect label information for training and evaluating protein secondary structure prediction models. To this end, we redesign the procedure for extracting primary sequence and secondary structure sequence information in this work. Specifically, for a given protein chain, we first extract its primary sequence from the sequence file pdb_seqres.txt.gz ^4^, which is provided on the PDB website according to the protein chains PDB ID and chain ID. For the convenience of distinction, we call this primary sequence the target primary sequence. Then, we obtain the corresponding dssp file according to the PDB ID. Then, contiguous amino acid residue fragments and corresponding secondary structure fragments are extracted from the obtained .dssp file according to the chain break identifier and residue sequence number. We generally cannot extract the primary sequence that is exactly the same or the same length as the target primary sequence from the .dssp file for two reasons: (1) Residues at the same sequence position are resolved to other types of residues; i.e., these re-didues have sequence microheterogeneity. (2) The coordinates of some residues cannot be resolved in X-ray crystallography or NMR spectroscopy experiments due to crystallographic disorder or physical gaps. Therefore, to obtain the corresponding secondary structure sequence of the target primary sequence, we used the Gotoh global alignment algorithm in the BioPython package to generate the alignment of the target primary sequence with each extracted residue fragment. The parameters match score, mismatch score, target gap score, query open gap score, and query extend gap score of the Gotoh alignment algorithm are set to 1, 0, −10, −0.01, and 0, respectively. For all subsequences with consecutive matching lengths not less than 5 in the alignment, their corresponding secondary structure fragments are assigned to corresponding residue fragments in the target primary sequence. Finally, for all the target primary sequence residues that are not assigned a secondary structure type, their secondary structure types are regarded as unknown. In the secondary structure sequence, we denote this unknown state with the letter X. In particular, X will be set to −100 in the label matrix **Y**.

We use four publicly available test sets, namely, CB433, CASP12, CASP13 and CASP14 to evaluate the performance of the proposed method. The CB433 dataset (Yang et al., 2022a) contains 433 protein chains, which were derived from the widely used CB513 dataset by merging protein chains with the same PDB ID and chain ID and replacing obsolete protein chains with newly released ones. The latter three datasets, CASP12, CASP13 and CASP14, were used in (Yang et al., 2022a,b), which contain 47, 41 and 33 protein chains, respectively. In particular, redundancy between protein chains was removed from these three datasets by using a sequence identity cutoff of 25%. To construct the corresponding training and validation sets, we selected protein chains from proteins with known structures based on the PISCES server (Wang and Dunbrack, 2003). The maximum resolution, maximum Rvalue, minimum chain length and maximum chain length of the screening criteria parameters were set to 2.5, 1.0, 50 and 800, respectively. In addition, protein chains with chain breaks or missing residues were also included. To obtain datasets of different sizes, we submitted six requests to the PISCES server on June 23, 2022, while considering the above settings and six different maximum sequence percentage identity thresholds of 25, 30, 35, 40, 45 and 50. For the sequence identity thresholds of 25, 30, 35, 40, 45 and 50, the numbers of protein chains returned by the PISCES server were 11735, 16242, 20465, 23967, 26899 and 29176, respectively. Moreover, these datasets were further filtered to remove protein chains that had more than 25% sequence identity to any protein chain in the four test sets. The filtered six datasets are called PISCES_25, PISCES_30, PISCES_35, PISCES_40, PISCES_45 and PISCES_50, which contain 11074, 15204, 18946, 22014, 24478 and 26304 protein chains, respectively. Finally, we randomly selected 512 protein chains from each dataset as the validation set and the remaining protein chains as the training set. Therefore, the numbers of protein chains in the six training sets are 10562, 14692, 18434, 21502, 23966 and 25792, respectively.

In addition, we further construct a larger independent test set. Specifically, we first obtain PDB IDs of all protein structures determined by X-rays released in the PDB database from July 1, 2022, to October 31, 2022, with the constraints of resolution < 2.5 Å and R-free < 0.25. Then, we used the PISCES server to cull nonredundant protein chains from the obtained PDB IDs according to the pairwise percent sequence identity < 25% and the minimum chain length = 50. Finally, the PISCES server returns 634 protein chains, and we call this test set CB634. For dataset CB634, we construct its corresponding training data based on 29,176 protein chains previously obtained using the sequence percentage identity threshold of 50. We remove all protein chains with > 25% sequence identity to a protein chain in the test set. The resulting dataset contains 25,805 protein chains. We randomly selected 512 protein chains as the validation set and the remaining 25,293 proteins as the training set.

The secondary structure information in the datasets mentioned above are all derived from nine-class secondary structure assignment. However, existing secondary structure prediction models were trained on datasets constructed from eight-class secondary structure assignment. Therefore, to fairly compare the proposed model with existing secondary structure predictors, evaluation experiments need to be performed on the dataset constructed from the old eight-class assignment. Here, we use two widely used test sets, namely, TEST2016 and TEST2018 (Hanson et al., 2019), which contain 1213 and 250 protein chains, respectively. In particular, these test sets correspond to the same training and validation sets, where the training set contains 10029 protein chains and the validation set contains 983 protein chains. The length of the longest protein chain in the four datasets does not exceed 700. Moreover, we use the default primary and secondary structure sequence information of these four datasets.

### 3.2. Implementation details

Our implementation is built on top of an adapter-transformer (Pfeiffer et al.), which is an extension of HuggingFace’s transformer library and integrates adapters and other efficient finetuning methods for transformer-based PyTorch language models. All our experiments are performed on a single NVIDIA GeForce RTX 3090 Ti GPU with 32 GB of memory. We train the proposed models by using the AdamW optimizer with *β*_1_ = 0.9, *β*_2_ = 0.999, a batch size of 32, and a fixed random seed of 0. For lightweight fine-tuning, both the learning rate and weight decay are set to 0.001. For full fine-tuning, both the learning rate and weight decay are set to 1e-5. In particular, the weight decay corresponding to the parameters of all layer normalization layers is set to 0 in both fine-tuning cases. Moreover, early stopping with a patience of 5 nonincreasing epochs is adopted to avoid overfitting.

Fine-tuning pretrained large models usually requires a large GPU memory space. For protein secondary structure prediction, we are faced with the inability of the 32 GB GPU to accommodate the whole batch size during training. To address this problem, we use three strategies: (1) For each protein chain with a length greater than 512 in a training set, we divide protein chain equally into two subchains. The added EOS token is now included in the chain length. Specifically, let the primary sequence of the protein chain to be divided be *R*_1_*R*_2_… *R*_*L*-1_*R*_L_, where *L* denotes the length of the protein chain. After division, the lengths of the two subchains should be ⌊*L/2* ⌋ and *L* - ⌊*L*/2 ⌋. To avoid losing the contextual information of each residue in the subchains, we set the primary sequences of the two subchains as *R*_1_*R*_2_ … *R*_511_*R*_512_ and *R*_*L*-511_*R*_*L*-510_ … *R*_*L*-1_*R_L_*, respectively. For the subsequence *R*_⌊*L*/2 ⌋+1_*R*_⌊*L*/2 ⌋+2_ … *R*_511_*R*_512_ in the first subchain and the subsequence *R*_*L*-511_*R*_*L*-510_ … *R*_⌊*L*/2 ⌋-1_*R*_⌊*L*/2 ⌋_ in the second subchain, their corresponding secondary structure label values are set to −100 to avoid introducing duplicate label information. The divided subchains are treated as independent protein chains during training. We did not do this for the protein chains in the validation and test sets. (2) Automatic mixed precision training by using both single- and half-precision representations is enabled to save GPU memory space and speed up training. (3) To calculate the gradients based on the whole batch size, we use a gradient accumulation technique, which first iteratively calculates the gradients in smaller batches and then runs the optimization step after accumulating enough gradients. In this study, we set the gradient accumulation steps to 16 for full finetuning and 8 for lightweight fine-tuning; the numbers of these steps differ because full fine-tuning needs to save the gradients of all parameters and thus requires more GPU memory space.

As in (Yang et al., 2022b), we assign the secondary structure category corresponding to the most similar centroid to each amino acid residue. For a fair comparison with existing deep predictors, the per-residue accuracy and segment overlap measure (SOV) are reported as evaluation metrics. In particular, the two metrics can be naturally applied to 3-, 8- and 9-state predictions, and their detailed definitions of these metrics can be found in (Yang et al., 2022b). The proposed LIghtweight Fine-Tuning framework for Secondary Structure prediction is called LIFT_SS, and its source code and used datasets are available at https://github.com/fengtuan/LIFT_SS.

### 3.3. Ablation study

To investigate the performance of the proposed method under different scenarios, we perform comprehensive ablation studies on the CB433 test set. Among the seven lightweight finetuning methods (namely, Adapter, Compacter, Prefix-tuning, LoRA, (IA)^3^, MAM adapter, and UNIPELT), only (IA)^3^ does not involve additional hyperparameter tuning. Therefore, we use (IA)^3^ as the default lightweight fine-tuning method in our comparative experiments. Moreover, for the convenience of comparison, only the overall per-residue accuracies Q_9_ and Q_3_ are reported.

#### 3.3.1. Ablation study on different training data

When constructing training data for protein secondary structure prediction, many existing works (Hanson et al., 2019; Klausen et al., 2019; Wang et al., 2011; Yang et al., 2022b; Zhou and Troyanskaya) set the maximum pairwise sequence identity threshold between protein chains in the training data to 25% or 30%. However, these two thresholds are too small and will filter many protein chains with known structures. Therefore, to make full use of the protein data with known structures, we believe that the maximum sequence identity threshold between protein chains within the training data could be set higher under the condition that the training data are nonhomologous to the test data. To determine the appropriate threshold, we constructed six datasets, namely, PISCES_25, PISCES_30, PISCES_35, PISCES_40, PISCES_45 and PISCES_50, based on pairwise percent sequence identity thresholds of 25, 30, 35, 40, 45 and 50, respectively. The homology of these datasets with the CB433 test set was removed by a filtering operation with a sequence identity cutoff of 25%. The protein chains contained in the six datasets are 11074, 15204, 18946, 22014, 24478 and 26304, respectively. Obviously, the higher the sequence identity threshold, the more protein chains are included. In addition, each dataset was further randomly divided into a training set and a validation set containing 512 protein chains.

To investigate the impact of training data on the performance of the proposed method, we conduct a comparative experiment on the CB433 test set. In particular, for the purposes of comparison, we introduce a baseline model that freezes the encoder of ProtT5-XL-U50 and only updates the parameters of the prediction head and centroid vectors during training. Obviously, the baseline model is similar to DML_SS^embed^. They differ only in the use of different basic building blocks. The performance comparison between the proposed method and the baseline model on different training data for nine-class and three-class predictions are illustrated in Figs. 5 and 6, respectively. Obviously, LIFT_SS((IA)^3^) always outperforms the baseline model in all cases. In particular, when trained on the PISCES_50 dataset, the Q_9_ and Q_3_ accuracies of LIFT_SS((IA)^3^) on the CB433 test set are 0.85% and 0.66% higher, respectively, than those of the baseline model. This finding validates the effectiveness of the proposed lightweight fine-tuning method in protein secondary structure prediction. Moreover, the figures reveal that the overall accuracies of both nine-class and three-class predictions increase as more protein chains are available in the training data. Particularly, compared to the results trained on PISCES_25, LIFT_SS((IA)^3^) trained on PISCES_50 improves by 1.11% and 0.87% on the Q_9_ and Q_3_ accuracies, respectively. The baseline model also shows similar improvements on nine-class and three-class predictions. This finding confirms that making full use of protein data with known structures can indeed improve the prediction accuracy of the secondary structure. Unless otherwise specified, we use the dataset PISCES_50 as our default training data in the following experiments. Of course, further increasing the training data by adjusting the maximum sequence identity threshold between protein chains within the training data will not significantly improve prediction accuracy but will increase training time. For the baseline model, the highest Q_9_ and Q_3_ accuracies on the CB433 test set are obtained on the training data PISCES_45 and PISCES_40, respectively. However, for the proposed method LIFT_SS((IA)^3^), when the training data are changed from PISCES_25 to PISCES_50, the method obtains almost continuous performance improvement with the increase in the number of protein chains available in the training data. In particular, LIFT_SS((IA)^3^) achieves both the highest Q_9_ and Q_3_ test accuracies on the training data PISCES_50.

**Fig. 5.**
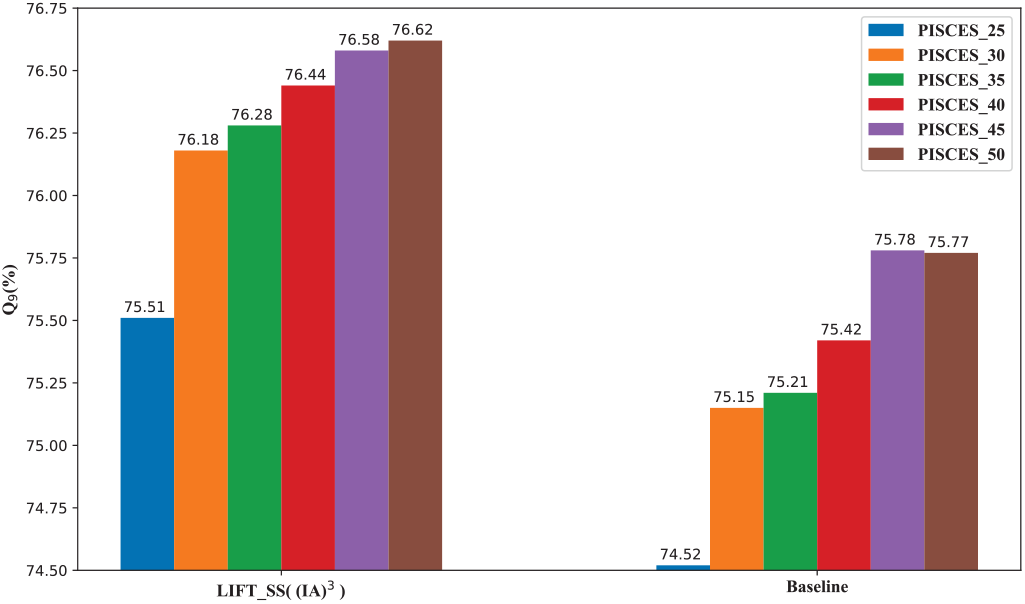
Comparison of nine-class secondary structure prediction results under different training data on the CB433 test set.

**Fig. 6.**
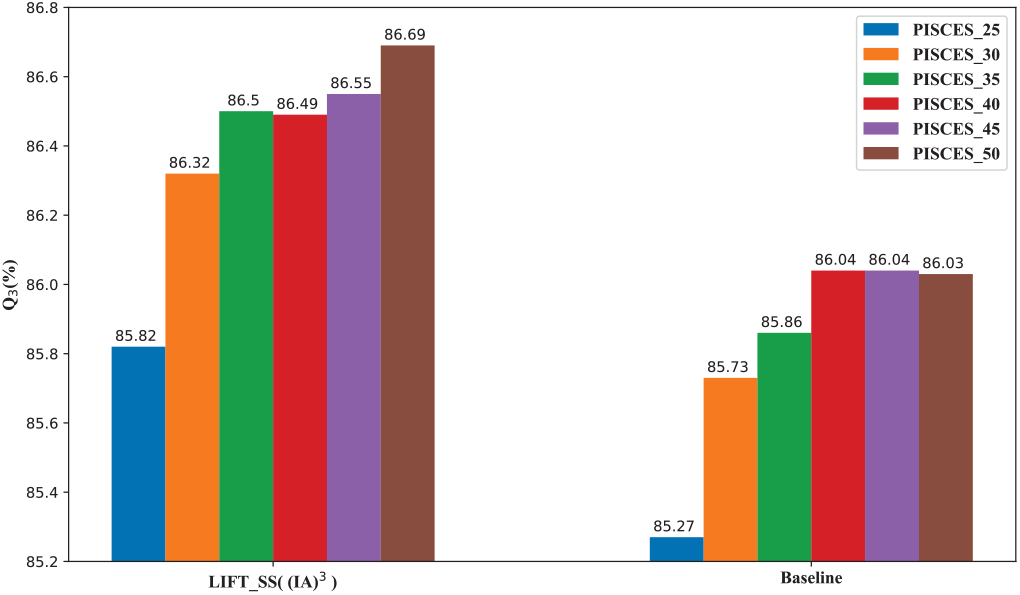
Comparison of three-class secondary structure prediction results under different training data on the CB433 test set.

#### 3.3.2. Ablation study on loss function

In our work, we adopt the softmax-based metric learning loss to train the proposed model. In fact, the commonly used cross-entropy classification loss can also be adopted when we replace the last normalize layer in the prediction head with a ReLU activation layer and a fully connected layer with c units and no bias. In both cases, the total number of parameters of the prediction head and loss function are the same. Hence, it is interesting to investigate the impact of both loss functions on the performance of the proposed method. Fig.7 shows the prediction results of the proposed method LIFT_SS((IA)^3^) under the two loss functions. As the figure indicates, the performance of the two loss functions on the CB433 test set is comparable, the cross-entropy loss slightly outperforms the metric learning loss in the nine-class prediction, and the latter slightly outperforms the former in the three-class prediction. In particular, we find that compared to the cross-entropy loss, the metric learning loss requires more training epochs to converge. This may be the reason why the metric learning loss achieves higher accuracy on the validation set.

**Fig. 7.**
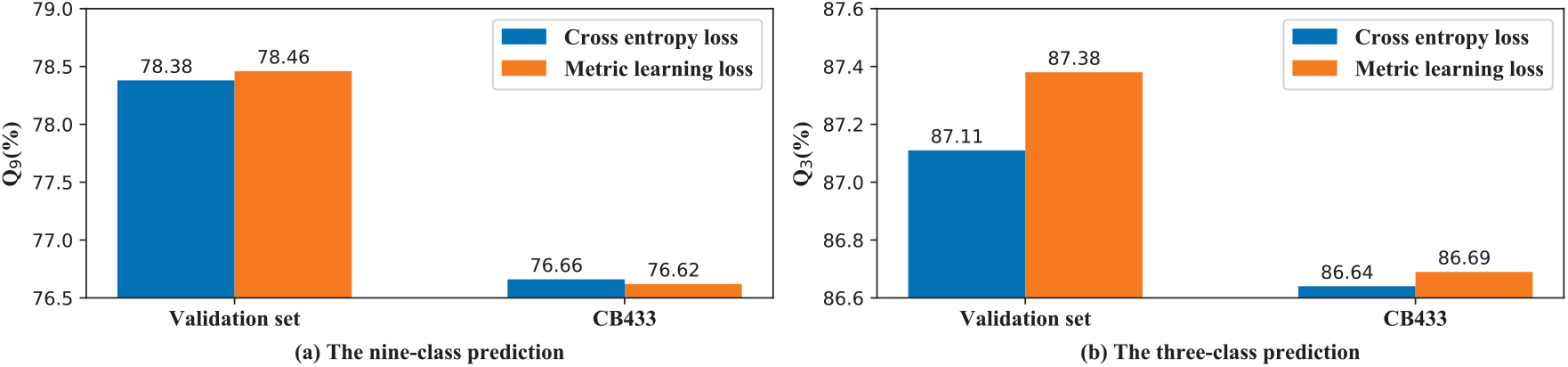
The prediction results of the proposed method LIFT_SS((IA)^3^) under the two loss functions.

#### 3.3.3. Ablation study on the fully fine-tuned model

##### Impact of learning rate

Based on the proposed secondary structure prediction framework, we can perform full model fine-tuning on the pretrained transformer model. In particular, the performance of the fully fine-tuned model significantly depends on the adopted learning rate. To determine the optimal learning rate, we compare the performance of the fully fine-tuned model on nine-class and three-class predictions at five different learning rates 1.0 × 10^-4^, 5.0 × 10^-5^, 1.0 × 10^-5^, 5.0 × 10^-6^, 1.0 × 10^-6^. The prediction results of the fully fine-tuned model under different learning rates are shown in Fig.8. The validation set is derived from the dataset PICSES_50. As the figure reveals, both the Q_9_ and Q_3_ accuracies of the fully fine-tuned model are the highest on the validation set when the learning rate is 1.0 × 10^-5^. Therefore, we use 1.0 × 10^-5^ as the default learning rate for full fine-tuning. Furthermore, we find that for smaller learning rates, the fully fine-tuned model requires more epochs to end the training.

**Fig. 8.**
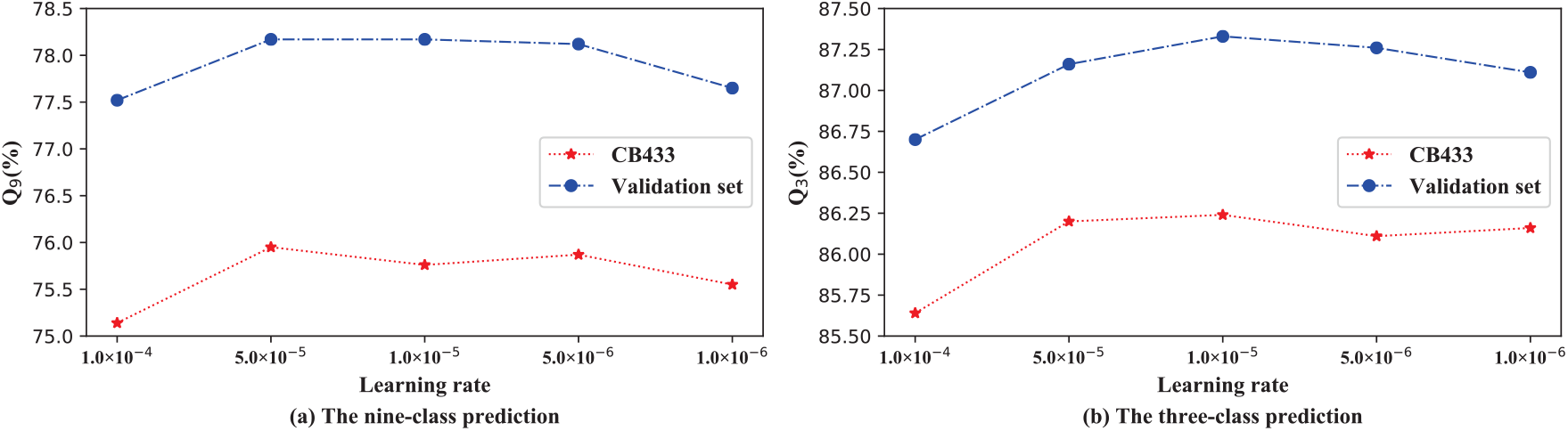
The prediction results of the fully fine-tuned model under different learning rates.

##### Impact of the initialization strategy

The initialization of the prediction head greatly affects the convergence of the fully fine-tuned model. In our experiments, the weights of the prediction head are initialized by the default random initialization strategy provided by PyTorch. However, we can also initialize the prediction head by using the weights of the trained baseline model. To train the baseline model, we can first save the embedding feature output by the encoder of ProtT5-XL-U50 and then update the parameters of the prediction head based on the loss function and the fixed embedding features. Therefore, the baseline model requires much less time than the fully fine-tuned model to complete training. In Table 1, we compare the impact of two initialization strategies on full model fine-tuning and lightweight fine-tuning. We also give the prediction results of the baseline model. As shown in Table 1, initializing the prediction head with the weights of the baseline model clearly improves the prediction performance of the fully fine-tuned model. In particular, the Q_9_ and Q_3_ prediction accuracies of the fully fine-tuned model are improved by 0.87% and 0.19%, respectively. However, a similar situation does not occur for the lightweight fine-tuning model LIFT_SS((IA)^3^). In contrast, the accuracies of Q_9_ and Q_3_ of LIFT_SS((IA)^3^) decreased, possibly because for the randomly initialized prediction head, the tuning of the pretrained model by full model fine-tuning at the beginning of training is wrong, thus causing the pretrained model to lose some learned useful information. Since the lost information cannot be recovered in subsequent optimization, the generalization performance of the fully fine-tuned model is poor. In contrast, lightweight finetuning can achieve better generalization performance by using random initialization because lightweight fine-tuning does not change the pretrained model. In particular, although initialization based on the baseline model can lead to faster training convergence, we find that lightweight fine-tuning offers only suboptimal performance. In the following, we discuss only the experimental results based on random initialization. As the table reveals, LIFT_SS((IA)^3^) outperforms the fully fine-tuned model by 0.86% and 0.45% in Q_9_ and Q_3_ accuracies, respectively. If the parameters of the prediction head and loss function are not considered, the latter requires adjusting approximately 1.4 billion parameters, while the former requires only adjusting 589,824 parameters. LIFT_SS((IA)^3^) is clearly more parameter-efficient. In our experiments, LIFT_SS((IA)^3^) and the fully fine-tuned model spend approximately 2200 s and 3300 s per training epoch, respectively. However, since the former requires more epochs to converge (e.g., in the nine-class prediction experiment, LIFT_SS((IA)^3^) runs 43 training epochs, while the latter only runs 17 epochs), the training cost of LIFT_SS((IA)^3^) is higher than that of the fully fine-tuned model.

**Table 1:**
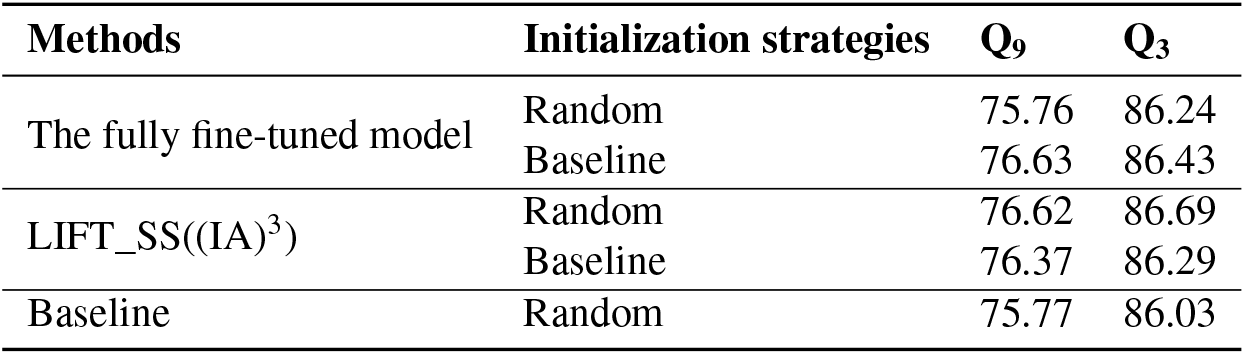
Comparison of the prediction results obtained by the two initialization strategies on the CB433 test set. “Random” indicates a random initialization strategy was used to initialize the prediction head. “Baseline” indicates that the weights of the trained baseline model were used to initialize the prediction head.

#### 3.3.4. Ablation study on the lightweight fine-tuning methods

For the proposed secondary structure prediction framework, we can use seven methods, namely, Adapter, Compacter, Prefix-tuning, LoRA, (IA)^3^, MAM adapter, and UNIPELT, to perform lightweight fine-tuning on the pretrained transformer model. To evaluate these lightweight fine-tuning methods, we conduct several experiments on the CB433 test set and its corresponding validation set derived from the dataset PICSES_50. Specifically, we analyze the impact of hyperparameters on the performance of the first four methods. (IA)^3^ does not introduce additional hyperparameters. The latter two hybrid methods also do not involve additional hyperparameter tuning once the hyperparameters of the first four methods are determined. For the lightweight fine-tuning methods, the default is to introduce new parameters for each transformer block. In fact, we can perform lightweight fine-tuning only for the top transformer blocks. In this way, there is no need to perform gradient calculations for lower transformer blocks. To this end, we further investigate the impact of the number of fine-tuned top transformer blocks on the prediction performance of the secondary structure. In this subsection, we report the experimental results only on the nine-class prediction.

##### Impact of bottleneck dimension

For Adapter and Compacter, the bottleneck dimension *d_bottleneck_* is an important configuration hyperparameter. Fig.9 shows the nine-class prediction results under the different bottleneck dimensions of 16, 32, 64, 128, and 256. On the CB433 test set, Compacter significantly outperforms Adapter. In particular, Compacter’s performance is not sensitive to changes in the bottleneck dimension.

**Fig. 9.**
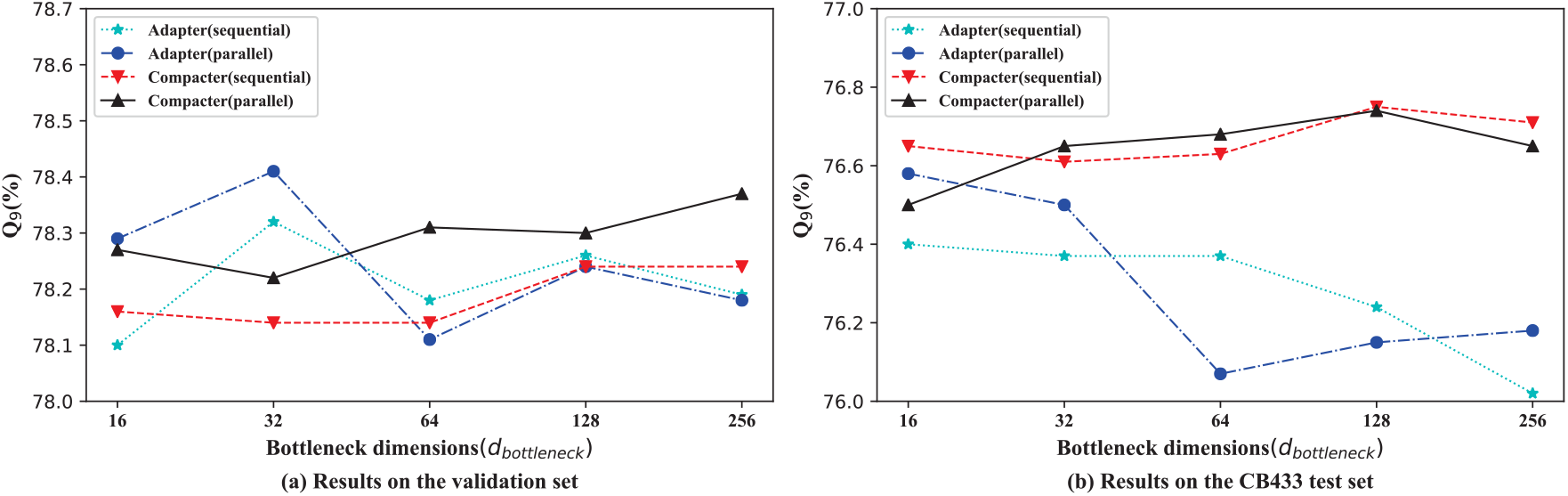
The nine-class prediction results of the proposed Adapter- or Compacter-based methods under different bottleneck dimensions.

However, Adapter’s performance tends to degrade as the bottleneck dimension increases possibly because the number of trainable Compacter parameters is almost constant across different bottleneck dimensions, while the number of Adapter parameters increases linearly with *d_bottleneck_*. Moreover, for both Adapter and Compacter, the parallel mode demonstrates no significant advantage over the sequential mode. This result differs from that in (He et al., 2022), possibly because we are targeting different tasks and different pretrained models. On the validation set, Compacter performs better in parallel mode than in sequential mode. Additionally, Compacter in parallel mode reaches the best validation accuracy at *d_bottleneck_* = 16. Hence, we use 16 as the default bottleneck dimension and consider only the parallel mode in the following experiments. In the two hybrid methods MAM adapter and UNIPELT, we use Compacter instead of Adapter since the former is more parameter efficient than the latter.

##### Impact of the prefix length

For the lightweight fine-tuning method prefix-tuning, the prefix length determines the number of trainable parameters to be added. As shown in Fig.10, the Q_9_ accuracy increases as the prefix length increases. When the prefix length is equal to 48, the accuracy on the validation set is the highest. Therefore, we use 48 as the default value for the prefix length. In addition, on the CB433 test set, the method’s accuracy is slightly lower when the prefix length is 48 rather than 40. Therefore, we did not increase the prefix length further in our experiments.

**Fig. 10.**
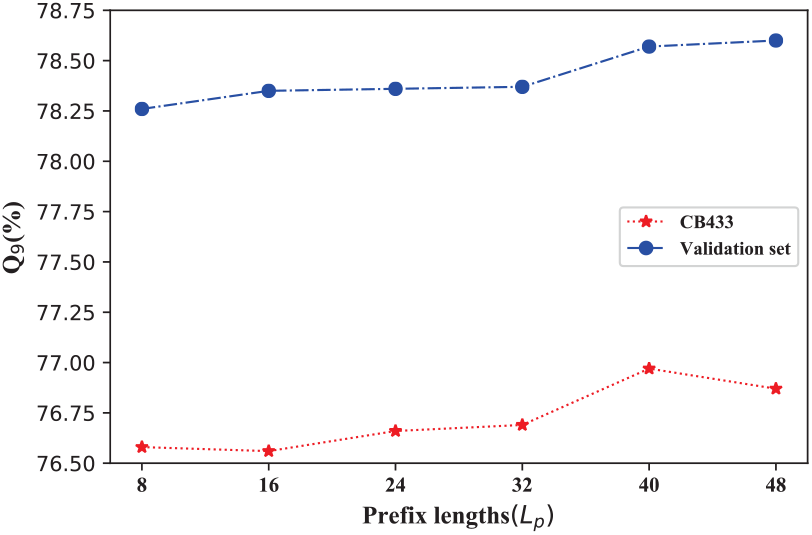
The nine-class prediction results of the proposed prefix-tuning-based method under different prefix lengths.

##### Impact of rank

We investigate the performance of the proposed LoRA-based method with varying ranks {1, 4, 8, 16, 32}. The prediction results are shown in Fig.11. On both the validation set and CB433 test set, the accuracy of Q_9_ decreases as the rank increases. This result suggests that when fine-tuning a pretrained protein language model, the update of the weight matrix has an extremely low “intrinsic rank”. In subsequent experiments, we set the rank *r* to 1.

**Fig. 11.**
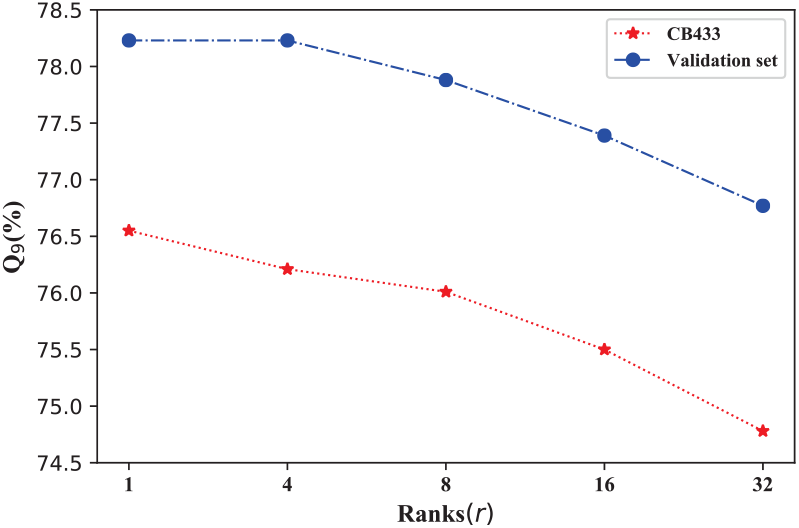
The nine-class prediction results of the proposed LoRA-based method under different ranks.

##### Impact of the number of fine-tuned top transformer blocks

For a pretrained model, we can fine-tune only the top transformer blocks and freeze the parameters of the other lower layers. Freezing these parameters eliminates the need to perform backpropagation on the lower layers, thus accelerating model training and reducing the amount of GPU memory required for training. Therefore, for lightweight fine-tuning in particular, we need to inject new parameters for only the top transformer blocks while keeping the other lower transformer blocks fixed. To investigate the impact of the number of fine-tuned top transformer blocks on the prediction performance of the seven lightweight fine-tuning methods, we conduct a comparative experiment on the CB433 test set and its corresponding validation set. The nine-class prediction results of the seven lightweight fine-tuning methods and full fine-tuning under different numbers of fine-tuned top transformer blocks are shown in Figs.12 and 13, respectively. The prediction accuracies of the seven lightweight fine-tuning methods on the CB433 test set are significantly better than those of the full fine-tuning in all cases. In particular, even when only the topmost transformer block is fine-tuned, the seven lightweight fine-tuning methods still demonstrate significant advantages although the parameters of the lower 23 transformer blocks are frozen at this point. A possible explanation is that full fine-tuning requires too many parameters to be adjusted, and therefore, its generalization performance is poor.

**Fig. 12.**
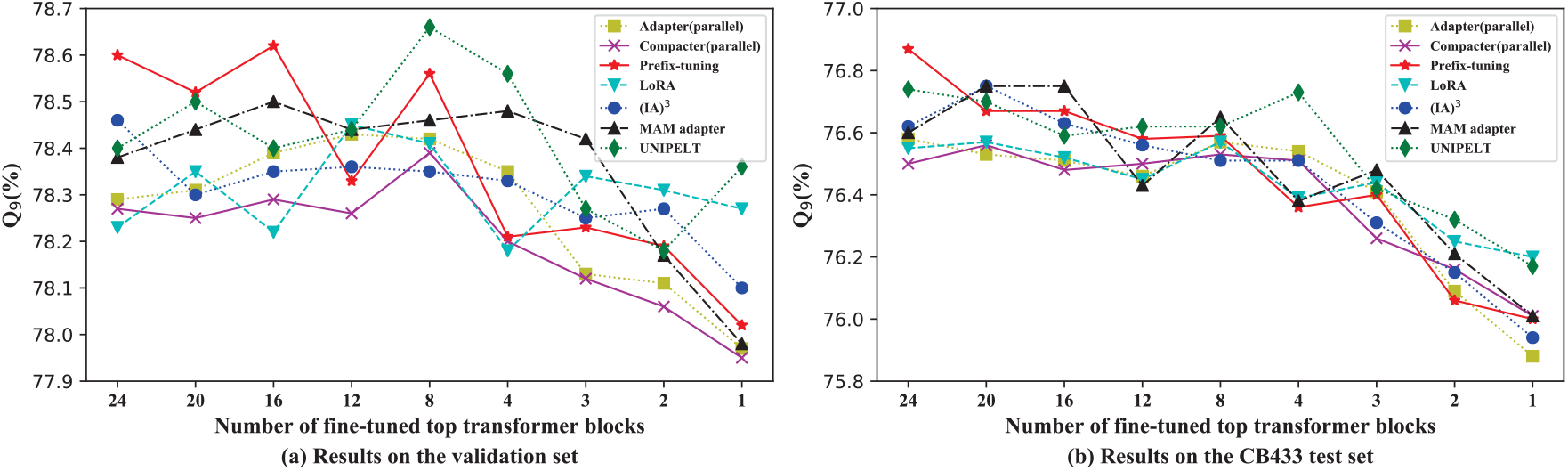
The nine-class prediction results of the seven lightweight fine-tuning methods under different numbers of fine-tuned top transformer blocks.

**Fig. 13.**
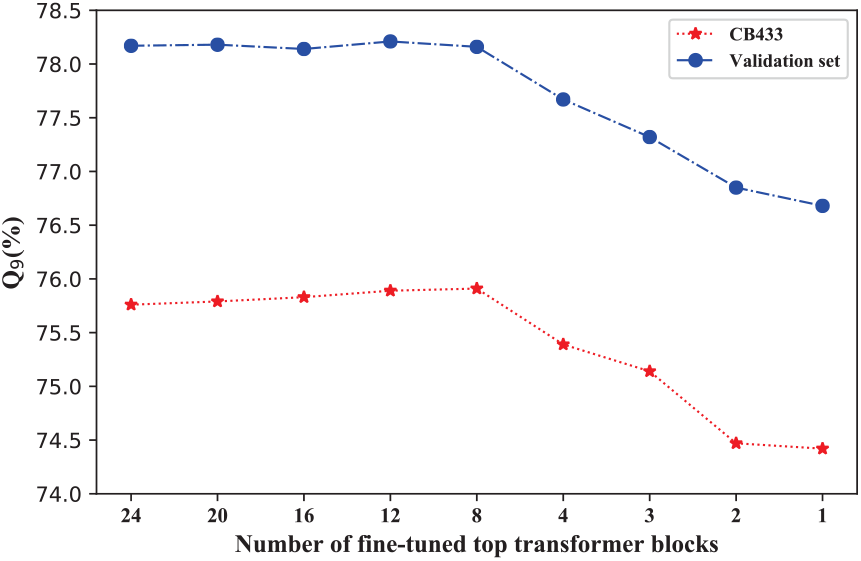
The nine-class prediction results of full fine-tuning under different numbers of fine-tuned top transformer blocks.

In addition, the performance of the seven lightweight methods on the validation and test sets fluctuates considerably when the number of fine-tuned top transformer blocks varies mainly because in the PyTorch implementation, the parameters of the network are randomly initialized sequentially from the lower layer to the higher layer. Although we use the same random seed, when a lower transformer block no longer introduces new parameters, the initial values of newly introduced parameters in the higher transformer blocks will change compared to the previous ones. Of course, the initial values of the parameters in the prediction head and loss function will change accordingly. More importantly, Fig.12 reveals that the prediction accuracies of all the lightweight fine-tuning methods on test set CB433 consistently fluctuate by approximately 76.6% with an amplitude of approximately 0.2% when the number of fine-tuned top transformer blocks is reduced from 24 to 8. This result suggests that in lightweight fine-tuning, only a few top transformer blocks need to introduce new parameters, while skipping many lower layers has little impact on the prediction accuracy of secondary structures. In particular, for the lightweight fine-tuning method LoRA, when the number of fine-tuned top transformer blocks is reduced from 24 to 8, the time spent per training epoch is also reduced from 1900 seconds to 1260 seconds. Training is accelerated by 33.7%. Inference will also accelerate correspondingly. Moreover, if we fine-tune all transformer blocks, the seven lightweight fine-tuning methods Adapter, Compacter, Prefix-tuning, LoRA, (IA)^3^, MAM adapter and UNIPELT need to introduce parameters of 3.5 MB, 0.56 MB, 36 MB, 1 MB, 2.27 MB, 38.3 MB, and 39.8 MB, respectively.The Compacter method clearly introduces the fewest parameters. When the number of fine-turned top transformer blocks is changed to 8, the number of parameters required for each lightweight finetuning method are reduced by two-thirds.

### 3.4. Comparison with state-of-the-art methods

In this section, we compare the performance of our proposed methods with existing state-of-the-art methods on two test sets, namely, TEST2016 and TEST2018. The compared methods include 10 profile-feature-based predictors (CNN_BIGRU (Drori et al., 2018), DeepACLSTM (Guo et al., 2019), DCRNN (Li and Yu), DeepCNN (Busia and Jaitly, 2017), MUFold-SS (Fang et al., 2018), NetSurfP-2.0 (Klausen et al., 2019), SPOD-1D (Hanson et al., 2019), SAINT (Uddin et al., 2020), SPIDER-3 (Heffernan et al., 2017), and DML_SS (Yang et al., 2022b)) and 2 embedding-feature-based predictors (SPOT-1D-LM (Singh et al., 2022b) and DML_SS^embed^ (Yang et al., 2022b)). The eight-class and three-class secondary structure prediction results of the proposed methods and the existing 12 predictors are given in Table 2. For a fair comparison, the results of the 12 existing predictors are taken from the published literature (Singh et al., 2022b; Yang et al., 2022b). The table reveals that the proposed three lightweight fine-tuning methods consistently outperform the other 12 predictors on the TEST2016 dataset. On the TEST2018 dataset, the proposed lightweight fine-tuning methods also perform better in most cases. The only exception is produced by DML_SS^embed^, which slightly outperforms the proposed lightweight fine-tuning methods on the SOV3 metric. These results demonstrate the effectiveness of our proposed lightweight fine-tuning framework. The sequence identity cutoff value between the protein chains in the training dataset corresponding to the two test sets TEST2016 and TEST2018 is 25%. If we construct the corresponding training set based on the 50% sequence identity cutoff and perform training, the prediction accuracy of the proposed lightweight fine-tuning methods on TEST2016 and TEST2018 will be further improved.

**Table 2:** Comparison with state-of-the-art methods on the TEST2016 and TEST2018 test sets. The best results are shown in boldface. The symbol “-” indicates that the result is unavailable.

| Methods | TEST2016 |  |  |  | TEST2018 |  |  |  |
| --- | --- | --- | --- | --- | --- | --- | --- | --- |
|  | Q<sub>8</sub> | SOV<sub>8</sub> | Q<sub>3</sub> | SOV<sub>3</sub> | Q<sub>8</sub> | SOV<sub>8</sub> | Q<sub>3</sub> | SOV<sub>3</sub> |
| CNN_BIGRU (Drori et al., 2018) | 73.91 | 70.92 | 85.04 | 81.61 | 72.78 | 68.75 | 84.17 | 79.41 |
| DeepACLSTM (Guo et al., 2019) | 75.19 | 73.67 | 85.62 | 82.6 | 73.42 | 71.32 | 84.66 | 80.05 |
| DCRNN (Li and Yu) | 72.19 | 68.63 | 83.72 | 78.39 | 70.6 | 65.82 | 82.75 | 75.1 |
| DeepCNN (Busia and Jaitly, 2017) | 74.54 | 71.56 | 85.14 | 79.31 | 72.75 | 69.18 | 84.16 | 76.83 |
| MUFold-SS (Fang et al., 2018) | 76.03 | 73.67 | 85.97 | 81.98 | 74.29 | 71 | 84.63 | 79.53 |
| NetSurfP-2.0 (Klausen et al., 2019) | - | - | - | - | 73.81 | 71.14 | 85.31 | 78.58 |
| SPOD-1D (Hanson et al., 2019) | 76.03 | 73.88 | 86.67 | 79.52 | 74.26 | 71.45 | 85.66 | 78.77 |
| SAINT (Uddin et al., 2020) | 76.23 | - | - | - | 74.48 | - | - | - |
| SPIDER-3 (Heffernan et al., 2017) | - | - | 84.66 | 75.62 | - | - | 83.84 | 73.89 |
| DML_SS (Yang et al., 2022b) | 76.62 | 74.6 | 86.1 | 82.72 | 74.82 | 72.23 | 84.83 | 80.5 |
| SPOT-1D-LM (Singh et al., 2022b) | - | - | - | - | 76.47 | - | 86.74 | - |
| DML_SS<sup>embed</sup> (Yang et al., 2022b) | 78.03 | 75.9 | 87.41 | 84.51 | 76.48 | 73.44 | 86.82 | **82.43** |
| LIFT_SS(Prefix-tuning) | 78.48 | 76.56 | 87.72 | 84.71 | 76.70 | 73.78 | 87.07 | 82.16 |
| LIFT_SS((IA)<sup>3</sup>) | **78.70** | **76.79** | 87.80 | 84.56 | 76.70 | **74.24** | **87.13** | 82.32 |
| LIFT_SS(Compacter) | 78.54 | 76.50 | **87.84** | **84.76** | **76.86** | 73.74 | 87.03 | 82.23 |

In addition, it is interesting to compare the two embeddingfeature-based methods SPOT-1D-LM and DML_SS^embed^. SPOT-1D-LM is an ensemble of 3 models with different network architectures, including 2-layer BiLSTM, multiscale-ResNet and a cascaded hybrid of the first two models. The input feature of SPOT-1D-LM is the concatenation of 20-dimensional one-hot amino acid encoding, the 1280-dimensional embedding feature extracted from the pretrained model ESM-1b and the 1024-dimensional embedding feature extracted from the encoder of the pretrained model ProtT5-XL-U50. In particular, SPOT-1D-LM is not trained based on the default training and validation sets of TEST2018. The training set used by SPOT-1D-LM consists of 38913 protein chains, and the maximum sequence identity cutoff between protein chains is 95%. To train the model, SPOT-1D-LM uses multitask learning to simultaneously learn both three- and eight-class secondary structure prediction, backbone torsion angle prediction, solvent accessibility prediction, half-sphere exposure prediction and contact number prediction. In contrast, DML_SS^embed^ is a very simple single-task learning model. The input feature of DML_SS^embed^ is only a 1024-dimensional embedding feature from the encoder of ProtT5-XL-U50, and the embedding network that DML_SS^embed^ uses is a lightweight multibranch convolutional network that does not use multiscale features. Moreover, DML_SS^embed^ is trained with the default training data of TEST2018, which contains 10029 protein chains. Although SPOT-1D-LM is more complex than DML_SS^embed^ in terms of network model, training manner and input feature, and although the number of protein chains contained in the training data it uses is more than three times that of the latter, its three- and eight-class prediction accuracies are still slightly lower than those of the latter. One possible reason is that the embedding feature from a pretrained model already contains the contextual information of amino acids, so the prediction accuracy of secondary structures cannot be improved by using a multiscale network architecture or a bidirectional LSTM that captures long-distance dependencies. Moreover, a disadvantage of SPOT-1D-LM is that it cannot predict protein chains longer than 1024, while DML_SS^embed^ has no such limitation. Another embedding-feature-based approach is NetSurfP-3.0 (Høie et al., 2022), which, like SPOT-1D-LM, also uses multitask learning to train the prediction model. Since the embedding feature used in NetSurfP-3.0 is extracted from the pretrained model ESM-1b, the prediction accuracies of its reported three- and eight-class secondary structures are significantly lower than those of SPOT-1D-LM. Therefore, NetSurfP-3.0 is inferior to DML_SS^embed^ in terms of prediction performance.

### 3.5. Experimental results on the five test sets

To investigate the performance of the proposed framework LIFT_SS under seven different lightweight fine-tuning methods, namely, Adapter, Compacter, LoRA, Prefix-tuning, (IA)^3^, MAM adapter and UNIPELT, we perform comparative experiments on five test sets, namely, CASP12, CASP13, CASP14, CB433 and CB634. The first four test sets correspond to the same training and validation sets, while the test set CB634 has its own training and validation sets. For comparison purposes, we also give the prediction results of DML_SS^embed^. The nine-class and three-class prediction results of the eight methods on the five test sets are given in Tables 3 and 4, respectively. The two tables reveal that the prediction accuracies of the seven lightweight fine-tuning methods are better than those of DML_SS^embed^ in most cases. In particular, all lightweight fine-tuning methods consistently outperform DML_SS^embed^ on two larger test sets, namely, CB433 and CB634. Moreover, a comparison among the seven lightweight fine-tuning methods reveals no clear winner among them. In particular, although the two hybrid methods LIFT_SS (MAM adapter) and LIFT_SS (UNIPELT) introduce more parameters for the pretrained model, these methods have no clear advantage over the other five methods in terms of secondary structure prediction accuracy.

**Table 3:** Comparison of nine-class secondary structure prediction results on the five test sets CASP12, CASP13, CASP14, CB433, and CB634. The best results are shown in boldface.

| Methods | CASP12 |  | CASP13 |  | CASP14 |  | CB433 |  | CB634 |  |
| --- | --- | --- | --- | --- | --- | --- | --- | --- | --- | --- |
|  | Q<sub>9</sub> | SOV<sub>9</sub> | Q<sub>9</sub> | SOV<sub>9</sub> | Q<sub>9</sub> | SOV<sub>9</sub> | Q<sub>9</sub> | SOV<sub>9</sub> | Q<sub>9</sub> | SOV<sub>9</sub> |
| DML_SS<sup>embed</sup> | 74.81 | 70.07 | 72.32 | 67.25 | 65.40 | 58.43 | 75.59 | 73.35 | 76.78 | 73.62 |
| LIFT_SS(Adapter) | **75.82** | **70.81** | **73.07** | **68.06** | 64.50 | 56.90 | 76.58 | 73.89 | 77.39 | 74.44 |
| LIFT_SS(Compacter) | 75.20 | 70.13 | 72.62 | 67.89 | **66.59** | **59.71** | 76.50 | 73.96 | 77.63 | **74.83** |
| LIFT_SS(LoRA) | 74.84 | 69.92 | 72.58 | 67.34 | 66.17 | 58.97 | 76.55 | 74.05 | 77.42 | 74.20 |
| LIFT_SS(Prefix-tuning) | 75.46 | 70.77 | 72.36 | 67.85 | 66.49 | 59.26 | **76.87** | **74.36** | 77.54 | 74.49 |
| LIFT_SS((IA)<sup>3</sup>) | 74.83 | 70.02 | 72.49 | 67.80 | 65.71 | 57.51 | 76.62 | 74.28 | 77.54 | 74.45 |
| LIFT_SS(MAM adapter) | 75.15 | 70.64 | 72.20 | 67.89 | 66.33 | 58.53 | 76.60 | 74.23 | 77.61 | 74.56 |
| LIFT_SS(UNIPELT) | 74.85 | 69.50 | 72.19 | 67.30 | 66.32 | 59.36 | 76.74 | 74.09 | **77.68** | 74.54 |

**Table 4:**
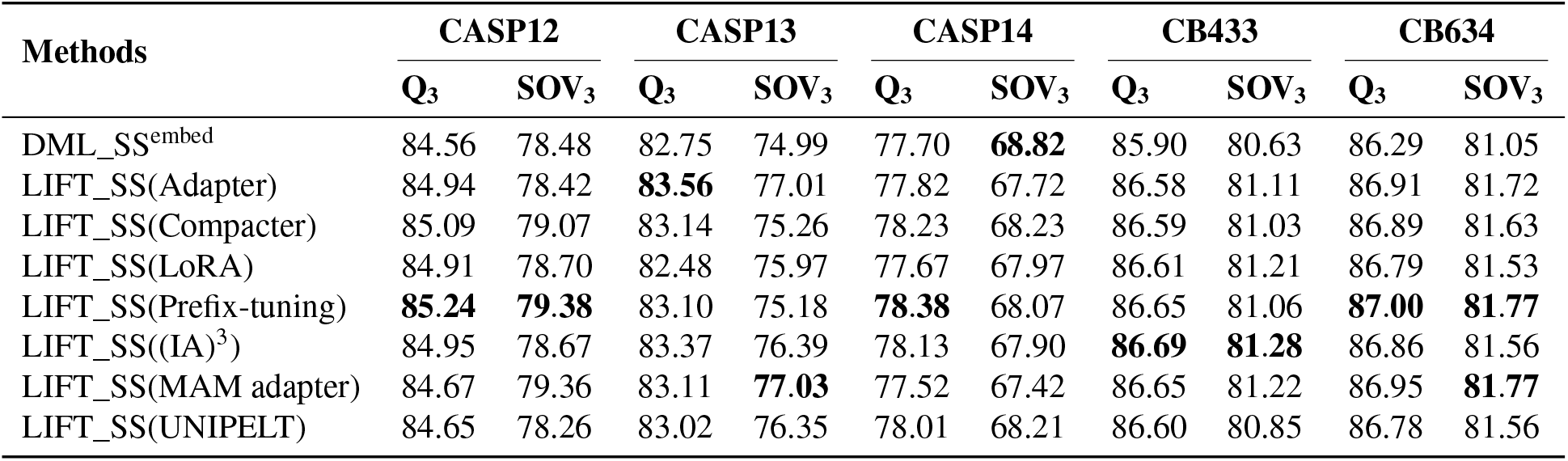
Comparison of three-class secondary structure prediction results on the five test sets CASP12, CASP13, CASP14, CB433, and CB634. The best results are shown in boldface.

### 3.6. Performance comparison of indirect and direct three-class prediction

For the nine-class predictor, the prediction probability sums can be calculated for 4 secondary structure types (H, G, I, and P), 2 secondary structure types (B and E), and 3 secondary structure types (T, L, and S). In this way, the predicted probabilities of coarse-grained 3-state secondary structure type helix, strand, and coil are obtained. In particular, we call this three-class prediction based on the nine-class predictor an indirect three-class prediction, and the prediction performed by directly training a three-class predictor a direct three-class prediction. On the basis of our proposed three methods LIFT_SS(Compacter), LIFT_SS(Prefix-tuning) and LIFT_SS((IA)^3^), we compare these two three-class predictions on two larger test sets CB433 and CB634. As shown in Table 5, the two methods do not have a clear winner in terms of prediction performance. Since the fine-grained nine-class prediction results and the coarse-grained three-class prediction results can be obtained with comparable performance simultaneously by the nine-class predictor, there is no need to train an additional three-class predictor for secondary structure prediction.

**Table 5:** Comparison of indirect three-class prediction and direct three-class prediction on the CB433 and CB634 test sets. The best results are shown in boldface.

| Methods | Type | CB433 |  |  |  |  | CB634 |  |  |  |  |
| --- | --- | --- | --- | --- | --- | --- | --- | --- | --- | --- | --- |
|  |  | Q<sub>C</sub> | Q<sub>E</sub> | Q<sub>H</sub> | Q<sub>3</sub> | SOV<sub>3</sub> | Q<sub>C</sub> | Q<sub>E</sub> | Q<sub>H</sub> | Q<sub>3</sub> | SOV<sub>3</sub> |
| LIFT_SS(Compacter) | 3 | 73.20 | 84.09 | 88.60 | 86.59 | 81.03 | 74.62 | 86.23 | 85.97 | 86.89 | 81.63 |
| LIFT_SS(Compacter) | 9 | 73.24 | 84.53 | 88.56 | 86.81 | 81.10 | 74.17 | 86.53 | 86.51 | 86.97 | 81.70 |
| LIFT_SS(Prefix-tuning) | 3 | **73.58** | 83.89 | 88.28 | 86.65 | 81.06 | **74.97** | 86.15 | 85.98 | **87.00** | **81.77** |
| LIFT_SS(Prefix-tuning) | 9 | 73.08 | 84.81 | 87.86 | **86.89** | 80.86 | 74.01 | 86.54 | **86.58** | 86.96 | 81.66 |
| LIFT_SS((IA)<sup>3</sup>) | 3 | 73.12 | **85.05** | **89.01** | 86.69 | **81.28** | 74.47 | **86.70** | 85.67 | 86.86 | 81.56 |
| LIFT_SS((IA)<sup>3</sup>) | 9 | 72.90 | 84.76 | 88.41 | 86.64 | 80.95 | 74.03 | 86.19 | 86.27 | 86.89 | 81.48 |

### 3.7. Ensemble prediction

Relying on the proposed method LIFT_SS(Compacter), we further investigate the performance of ensemble prediction on two tests CB433 and CB634. As in (Yang et al., 2022b), we train LIFT_SS (Compacter) according to five different random seeds 0, 10000, 20000, 30000 and 40000. The result of the ensemble prediction is the average of five individual model outputs. The prediction results of the ensemble model and its five individual models are given in Table 6. As the table reveals, the ensemble prediction clearly outperforms the best individual model. Moreover, each individual model requires only 0.56 MB to store the parameters of Compacter and 17.9 MB to store the parameters of the prediction head and loss function. Considering that the size of the encoder of the pretrained model ProT5-XL-U50 is 4.5 GB, the effect of ensemble prediction on the size of the secondary structure prediction model is negligible. In particular, if full model fine-tuning is used for ensemble prediction, we need to store five copies of the pretrained model. Thus, full model fine-tuning clearly results in higher storage and deployment costs. Therefore, lightweight fine-tuning is superior to full-model fine-tuning in terms of ensemble prediction.

**Table 6:** The nine-class prediction results of the ensemble model and its five individual models on the CB433 and CB634 test sets. The best results are shown in boldface.

| Methods | Seeds | CB433 |  | CB634 |  |
| --- | --- | --- | --- | --- | --- |
|  |  | Q<sub>9</sub> | SOV<sub>9</sub> | Q<sub>9</sub> | SOV<sub>9</sub> |
| LIFT_SS(Compacter) | 0 | 76.50 | 73.96 | 77.63 | 74.83 |
| LIFT_SS(Compacter) | 10000 | 76.84 | 74.23 | 77.52 | 74.53 |
| LIFT_SS(Compacter) | 20000 | 76.65 | 74.13 | 77.61 | 74.52 |
| LIFT_SS(Compacter) | 30000 | 76.48 | 73.96 | 77.54 | 74.50 |
| LIFT_SS(Compacter) | 40000 | 76.61 | 74.03 | 77.67 | 74.68 |
| **Ensemble model** |  | **77.27** | **74.68** | **78.13** | **75.13** |

## 4. Conclusion

In this paper, we propose a novel end-to-end protein secondary structure prediction framework called LIFT_SS, which learns task-specific representations by performing lightweight fine-tuning on a pretrained protein language model and thus significantly improves the performance of both three- and nine-class secondary structure prediction. In the proposed framework, each transformer block in the pretrained model is injected with a few new parameters. During training, we update only the newly introduced parameters, the parameters of the prediction head and the parameters of the loss function while keeping the original parameters of the pretrained model unchanged. This lightweight fine-tuning strategy not only significantly reduces the computational and storage overhead during training but also outperforms full model fine-tuning in predicting secondary structures. Moreover, we further improve the prediction accuracy of the secondary structure by ensembling five individual models trained with different random seeds. Extensive experimental results on seven benchmark test sets verify the effectiveness of the proposed framework. To our knowledge, this is the first work developed for nine-class secondary structure prediction. We experimentally show that the nine-class predictor can provide coarse-grained three-class prediction results comparable to those of the three-class predictor; this finding indicates that it is not necessary to train the three-class predictors separately. More importantly, our comparative experiments also demonstrate that simply increasing the sequence identity cutoff between protein chains can increase the size of the training data and thus significantly improve the prediction accuracy of secondary structures. In a broad sense, our research underpins the importance of lightweight fine-tuning in downstream applications of pretrained protein language models. In future work, we plan to apply lightweight fine-tuning techniques to other biological modeling tasks such as subcellular localization and fold recognition.

## Acknowledgement

This work was supported by the National Science Foundation of China (Grant no.61806074).

## Footnotes

1 https://github.com/PDB-REDO/dssp

2 ftp://ftp.cmbi.umcn.nl//pub/molbio/data/dssp/

3 https://cdn.rcsb.org/etl/kabschSander/ss.txt.gz

4 https://ftp.wwpdb.org/pub/pdb/derived_data/pdb_seqres.txt.gz

